# Evaluating the association between vocalizations and colouration in finches (Fringillidae), a passerine group with elaborate signals

**DOI:** 10.64898/2025.12.02.691818

**Authors:** A.I. Casale, P.L. Tubaro, D.A Lijtmaer

## Abstract

The interaction between multiple secondary sexual characters simultaneously expressed by an organism remains poorly understood. In fact, different interpretations of the role of concurrently expressed signals predict that the correlation of their elaboration across species should be positive, negative or absent —all three possibilities have been supported by studies of different avian groups. In this context, we analysed the interaction between plumage and song elaboration in the passerine family Fringillidae (finches). We used a two-scale approach and studied this association both at the family level and in more detail within the genera *Crithagra* and *Spinus*. At both scales, we determined colour elaboration (defined by the number of plumage colour patches in each species and their contrast) and established its association with three aspects of vocal elaboration: energy investment (song length and rate of syllable production), song complexity (repertoire index) and vocal performance (vocal deviation). We found an absence of association between colour elaboration and the three evaluated aspects of song elaboration at both scales, which has been the most frequent result in other avian groups. At first glance, this result appears to be partially discordant with previous findings in finches, but considering these studies together this might not necessarily be the case. We also confirmed that finches have complex songs and a remarkable vocal performance compared to other families of passerines. Considered together, our results suggest an independent evolution of colour and the notable vocal elaboration present in finches.

## Introduction

Since the foundational work of Darwin (1871), the evolution of secondary sexual traits has been a key component of evolutionary biology. This generated a vast amount of research, including contrasting hypotheses and debates on the mechanisms behind the evolution of these traits (Fisher 1930, Zahavi 1975, Prum 2012), the development and function of female ornamentations (Clutton-Brock 2007, Tobias *et al*. 2012), and the evolution of mate choice (Andersson & Simmons 2006, Kokko *et al*. 2006). A particularly debated aspect is the evolution of multiple, simultaneously expressed secondary sexual traits, a relatively understudied process despite the frequent occurrence of multiple ornaments in various animal groups (Møller & Pomiankowski 1993, Hebets & Papaj 2005). Most studies have focused on analysing single traits, without considering the presence of other sexual characters, yet understanding the interactions between these simultaneously expressed traits is crucial for a comprehensive understanding of ornament evolution. This perspective is particularly relevant for exploring behaviour, adaptive radiation, and speciation (Ligon *et al*. 2018).

From a theoretical point of view, three possible scenarios could be expected in terms of the interaction between secondary sexual characters: (1) If selection operates in the same direction on different characters, their complexity is expected to correlate positively. This could occur if each ornament conveys information about a particular property of the overall quality or condition of an individual (multiple message hypothesis), or if there is redundancy in the transmission of the message (redundant signal hypothesis) (Møller & Pomiankowski 1993, Johnstone 1996). (2) By contrast, Darwin himself (Darwin 1871) postulated that birds with more elaborate vocalizations usually have less elaborate colours, suggesting a negative correlation between the two types of signals, an idea that was later formalized as the ‘transfer hypothesis’ (Gilliard 1956). This hypothesis states that, because two costly signals that convey the same message would render an unnecessary investment, a signal emitted by a secondary sexual character should be replaced by a second signal that emerges, as long as the new signal is less expensive and capable of fulfilling the role of the pre-existing signal. In an intermediate scenario in which both signals are present and are costly, a negative correlation in the elaboration of the characters would be expected due to a trade-off in the investment in each of them (Shutler 2011). The cost in this context can be related to the energetic expenditure in signal production or other types of costs, such as an increased risk of predation. (3) Finally, if the multiple traits evolve independently, no correlation is expected among them. This could occur, for instance, if these signals are under different selection forces, as it would happen for example if they are associated with different ecological or life-history traits (Gomes *et al*. 2017) or if some of them are targeted by intersexual selection and others by intrasexual selection (Shutler & Weatherhead 1990).

The interest in understanding the interplay among secondary sexual characters increased in the last couple of decades, with studies across diverse animal groups (e.g., Kekäläinen *et al*. 2010, Hebets *et al*. 2013, Higham *et al*. 2013, Fichtel & Kappeler 2022, Webster *et al*. 2023), and with a particular focus in birds (e.g., Badyaev *et al*. 2002, Gonzalez-Voyer *et al*. 2013, Mason *et al*. 2014, Beco *et al*. 2021, Cardoso *et al*. 2024). Most of these avian studies have concentrated on the relationship between acoustic and visual signals. There are various reasons for this. In the first place, they constitute the principal channels of communication in birds, which often combine elaborate plumage and songs (Barreira & García 2019). In fact, these are the signals considered to be most influenced by sexual selection in birds (Darwin 1871, Catchpole 1987). Both signals are in addition considered to be expensive in terms of the energy investment that they require (Møller 1996, Oberweger & Goller 2001, Ward *et al*. 2003), a relevant aspect to consider for their study in the context of sexual selection, and in particular when analysing simultaneous signalling. On top of their energetic cost, conspicuous plumage colouration and singing can both have a cost associated with an increased predation risk (Zuk & Kolluru 1998, Huhta *et al*. 2003, Schmidt & Belinsky 2013, Bliard *et al*. 2020; but see Cain *et al*. 2019).

The increasing number of studies exploring the association between plumage and song elaboration has yielded contrasting results. Some analyses reported negative correlations, supporting the transfer hypothesis, including the study of cardueline finches (Badyaev *et al*. 2002), antwrens (Beco *et al*. 2021) and tits and chickadees (Tietze & Hahn 2024). However, other studies showed the opposite pattern, evidencing a positive correlation between the complexities of plumage colour and acoustic signaling in birds of paradise (Ligon *et al*. 2018), woodpeckers (Cárdenas-Posada & Fuxjager 2022) and canaries, goldfinches and allies (Cardoso *et al*. 2024). Finally, most studies did not find a correlation between characters, including the analysis of trogons (Ornelas *et al*. 2009), Asian barbets (Gonzalez-Voyer *et al*. 2013), tanagers (Mason *et al*. 2014), estrildid finches (Gomes *et al*. 2017), orioles (Matysioková *et al*. 2017), blue cardinalids (Barreira & García 2019) and parrots (Marcolin *et al*. 2022). This apparent inconsistency among studies could stem from their different methodological approaches (including the quantification of signal elaboration or complexity) or from biological disparities among avian groups, such as different types and degrees of selection affecting their characters or the different costs of the signals that they employ (Mason *et al*. 2014, Barreira & García 2019).

Fringillidae is a particularly suitable passerine family for studying the relationship between plumage colouration and vocal signals, given the marked variation among its members in both colour and song elaboration, including in particular species with highly complex vocalizations (Cardoso & Mota 2007, Funghi *et al*. 2015, Cardoso *et al*. 2020). In addition, both plumage colouration and song complexity have been shown to be under sexual selection, as evidenced by female preferences for more elaborate traits in both modalities (Hill 1990, Cardoso *et al*. 2012). Finally, previous studies performed on finches have shown contrasting results, including both negative (Badyaev *et al*. 2002) and positive (Cardoso *et al*. 2024) correlations between the complexity of these signals. In this context, we examined the interaction between male plumage colouration and vocalizations at two different taxonomic scales: (1) at the family level, by comparing the songs between the species with the most and the least elaborate colour pattern in each finch genus and (2) within two genera, *Crithagra* and *Spinus*. These two genera were chosen because their members vary gradually in their colouration and, in the case of *Spinus*, because it includes some of the species with the most complex songs in the family. In addition, *Spinus* was one of the genera that was analysed both by Badyaev *et al*. (2002) and Cardoso *et al*. (2024), making it an interesting case for comparison with their results. At both scales, we studied the association between plumage elaboration and three aspects of vocal elaboration: energy investment, song complexity and vocal performance.

## Methods

### Plumage colour elaboration analysis

We based the assessment of colour elaboration on the number of colour patches and their contrast, as these are considered key components of avian colour pattern advertising (Endler 1990, 1992). These variables have been widely used to study the evolution of colour elaboration and display in birds (Heindl & Winkler 2003, Gomes *et al*. 2017, Beco *et al*. 2021) and other animals (Pérez i de Lanuza *et al*. 2016, Kemp *et al*. 2023). Colour elaboration was evaluated for all extant finch species that were represented either in illustrations in the Handbook of the Birds of the World (Collar *et al*. 2010) or in photographs from the Macaulay Library (http://macaulaylibrary.org/). Although book illustrations or pictures are contentious for the assessment of hue or brightness in specific plumage patches, they are a robust source for analysing colour patterning, especially in studies with broad taxonomic and geographic scales that complicate the use of museum collections (e.g., Martin *et al*. 2010, Somveille *et al*. 2016).

For each species, we identified and counted all colour patches present in male plumage, regardless of female colouration. Patches were defined as continuous body areas of the same colouration. Areas exhibiting streaked or spotted patterns were considered a single patch, regardless of the presence of more than one colour. We then scored the contrast between each pair of adjacent patches using three levels: high contrast = 2, medium contrast = 1, and low contrast = 0.5. Based on these data, we generated a Colour Elaboration Index (CEI) that integrated both the number of patches and the contrast between them. The CEI for each species was calculated as the number of plumage patches multiplied by the mean of the contrast scores of all pairs of adjacent patches (i.e., CEI values increased with more patches and higher contrast among them). In cases where there were illustrations or pictures from more than one subspecies, we averaged the CEI values across subspecies to obtain a single final value per species. The CEI was calculated for all species by the same researcher (AC) prior to song analyses, ensuring that the classification was blind to vocal elaboration.

Taxonomy followed the checklist by Clements *et al*. (2024), and colour elaboration was evaluated for a total of 208 extant species belonging to 42 genera of Fringillidae (7 genera were excluded because they do not include any extant species). The list of these species and their CEI values is reported in Table S1.

### Song elaboration analysis

We downloaded all available male recordings from the selected species at the family-scale analysis (see the following section for the description of the criteria to select the species) and all recordings from *Crithagra* and *Spinus* species from the Macaulay Library and the online repository Xeno-Canto (http://xeno-canto.org/). At both taxonomic scales, we analysed a maximum of ten individuals per species (depending on availability). When recordings from more than ten individuals were available, we chose ten high-quality recordings choosing localities that allowed maximizing the geographic representation of the species. Details of the number of individuals per species included in each analysis and their corresponding recording catalogue numbers are provided in Tables S2-S4. For the family-scale analysis and for the genus *Crithagra* we used three songs per individual (except in cases where fewer than three songs were available). For the genus *Spinus*, we randomly selected ten-second fragments for the analysis due to the extended duration of the songs, which precluded conducting the analysis of the entire song (song length was not measured for this analysis; see below). To select these fragments, a random number generator was used to choose starting points within the recordings, ensuring that each selected fragment included at least 30 seconds of audio beyond the starting point and had an adequate signal-to-noise ratio. A total of 1248 songs were analysed from the 92 species of the data set (817 songs at the family level, 423 *Crithagra* songs and 80 *Spinus* songs; 72 songs were used both at the family and the genus levels, as they belonged to the species with the most and the least elaborate plumage colour in *Crithagra* and *Spinus*).

We used Raven Pro v. 1.5 (K. Lisa Yang Center for Conservation Bioacoustics, Ithaca, NY, USA) to visualize and measure songs. For standardization of measurements, all spectrograms were generated using a Hann window with 300 samples, a DFT size of 512 samples and the “cool” colour scheme, keeping brightness and contrast at 50%.

We considered the syllable as the basic unit of the song, defined as a single element or a set of elements produced with a minimum temporal separation (Cardoso & Mota 2007, Catchpole & Slater 2008). The repetition of a syllable with a simple structure was considered a trill. Finally, we considered a succession of syllables separated by a considerable period of silence with respect to the previous and subsequent succession as an individual song.

We measured 8 temporal- and frequency-related variables for each song: song length, trill length, maximum frequency, minimum frequency, bandwidth (the difference between the maximum and the minimum frequencies), number of syllable types, total number of syllables and syllable rate. The first 5 variables were measured by selecting each song in the spectrogram and using the corresponding Raven Pro measurement options (Fig. 1). The number of syllable types was determined by visually identifying all syllables in the spectrogram and classifying them into types according to their overall temporal and frequency structure (i.e., syllables with similar contour shape and duration were considered the same type; Catchpole & Slater 2008). The total number of syllables was determined by counting the syllables in the spectrogram and the syllable rate was quantified by dividing the total number of syllables by song length (thus obtaining the number of syllables emitted per second; Podos 1997). The measured song variables were used to study three aspects of vocal elaboration: energy investment, song complexity and vocal performance (Podos 1997, 2001, Cardoso & Mota 2007, Podos 2009, Cardoso & Hu 2011). The energy investment of males was analysed considering two of the variables described above: song length and syllable rate. The former informs the time spent on each song, and the latter considers the proportion of that time in which the bird is actually producing sound. Both are standard variables used in song analysis (Catchpole & Slater 2008). Both variables were calculated for each song, then averaged among songs to obtain a value for each individual, and finally averaged among all the individuals of each species.

**Figure 1.**
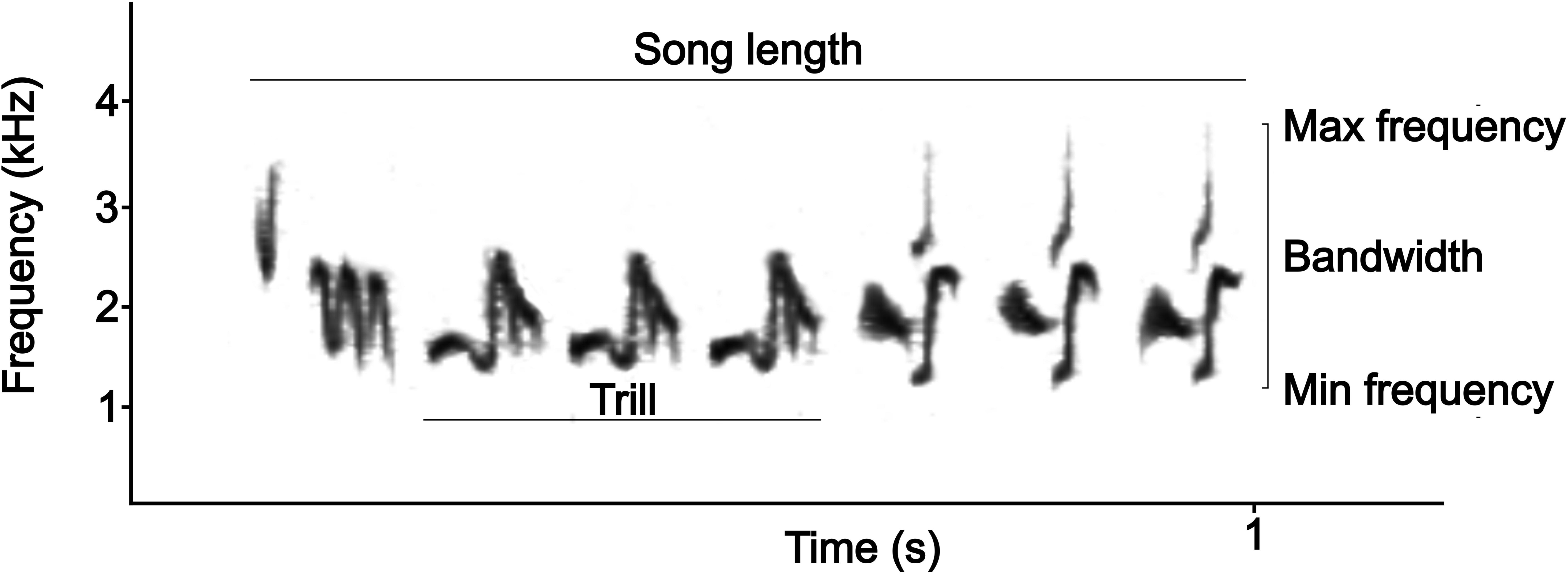
Scheme of the variables measured in Raven Pro v. 1.5, using a fragment of a *Spinus magellanicus* song as an example. The five variables shown in the figure were measured directly on the spectrogram using the Raven Pro measurement options. The number of syllable types was determined by identifying in the spectrogram all the syllables used in the song (3 in this scheme), the total number of syllables by direct counting (8 in this scheme), and the syllable rate by dividing the total number of syllables by song length (8 syllables / 0.9 s = 8.89 Hz in this scheme).

Song complexity was estimated by calculating the repertoire index (Cardoso & Mota 2007), which considers the degree of differentiation between the syllables of successive songs (Bell *et al*. 2004, Garamszegi *et al*. 2005, Cardoso & Mota 2007). This variable has the advantage of allowing the study of complexity without the need to analyse the entire repertoire of each individual, which is not possible in most cases and certainly unrealistic when working with recordings from sound libraries. Following Cardoso & Mota (2007), for each pair of successive songs we calculated the repertoire index as the average number of syllable types per song, multiplied by 1 + (1 – the proportion of shared syllable types) (if the number of syllable types differed between songs, the proportion was calculated using the song with fewer syllables). The higher the index, the more complex the song is (because a lower proportion of syllables is repeated between consecutive songs). In fact, this index can vary between a minimum value equivalent to the number of syllable types of each song (if the songs share all syllable types) and a maximum value equivalent to the sum of the syllable types of both songs (if no syllable type is shared). This index was calculated three times for each individual using three songs (three paired comparisons among them) and we then averaged them to obtain the repertoire index of the individual (if only two songs were available from an individual, we calculated the index only once, and this variable was not calculated for individuals with only one recorded song). In the case of the analysis within the *Spinus* genus, three 10-second fragments were used from each individual instead of three complete songs, as for the rest of the variables. Finally, the repertoire indexes of all the individuals of each species were averaged to obtain the species value.

We evaluated vocal performance as the third aspect of song elaboration because it has been proposed that males could demonstrate their quality and attract females not only through complex songs, but also by the presence of simple structures (trills) that allow forcing the male’s phonation system close to its physiological limit (Podos 2009, Cardoso & Hu 2011; see Irschick *et al*. 2008, for a review about selection operating on performance capabilities). This limitation is associated with the maximum bandwidth a bird can utilize at a given trill syllable rate. In essence, because trills typically involve rapid frequency modulations, producing fast trills with brief intervals between syllables imposes physiological constraints on the vocal apparatus. At least two distinct mechanisms underlie this constraint. First, the frequency of a syllable at any given moment is determined by the tension of the syringeal labia, which itself is controlled by the contraction level of the syringealis ventralis muscles (Laje *et al*. 2002). Consequently, the muscle’s maximum contraction and relaxation speed restrict the bandwidth achievable when syllable intervals become very short (Lijtmaer 2008). In addition, tonal frequency also depends on the configuration of the vocal tract, including the trachea, mouth and beak. Since these structures also have limits in their movement speed, faster trills inevitably result in reduced bandwidth, a pattern demonstrated in a large-scale study of Emberizidae (Podos 1997).

The mechanism of sound production differs between relatively slow trills and very fast, buzz-like trills, both in terms of their respiratory patterns and the movements of the syrinx and the upper vocal tract. First, in slower trills syllables are generated during expirations, with rapid inspirations occurring between them, but in fast trills with very short intervals between syllables, these mini-breaths are replaced by a pulsatile pattern, in which bronchial pressure is high throughout the trill, without inspirations, and the air passage across the syrinx is actively controlled by its muscles (Hartley & Suthers 1989, Hartley 1990, Mindlin & Laje 2005, Lijtmaer 2008). Second, both the syrinx and the vocal tract behave differently between slow and fast trills, as the cycles of bronchial air pressure and syrinx muscle tension that determine the characteristics of each syllable are clearly modified (Mindlin & Laje 2005, Lijtmaer 2008), as are the vocal tract cyclic movements when the bird is singing (Podos 1997). As a consequence of these differences in sound production, very fast trills are not necessarily limited in their bandwidth as much as the slower trills (i.e., an increase in their syllable rate might not necessarily compromise their bandwidth) and are therefore not as appropriate to study vocal performance. Even though the exact limit between these two modes of song production is not known, and it might actually differ among bird groups or species, we followed Podos (1997) and considered only relatively slow trills (i.e., excluding trills with rates over 50 Hz) for vocal performance analysis.

We assessed vocal performance by measuring vocal deviation (Podos 1997, 2001), which indicates how much each particular trill (given its rate) deviates from the maximum physiological performance (maximum bandwidth) that is defined for the entire Fringillidae family. To calculate this, we first measured trill rates in the same way as syllable rates, but considering each trill separately and dividing the total number of syllables in each trill by its length. We then plotted bandwidth as a function of trill rate using all trills in the vocal dataset with rates not higher than 50 Hz. After this, we split trill rate values into 4 Hz windows and selected the trill with the highest bandwidth in each of them. An upper-bound regression using these selected trills was established as an indicator of the maximum physiological performance as a function of trill rate at the family level. We calculated the vocal deviation of each trill as the orthogonal distance (in Hz) to this upper-bound regression using the *dplyr* package in R (Wickham 2015), then averaged all trills of an individual to obtain its performance value and finally averaged all individuals from each species. This allowed us establishing how much each species forces its phonation system when singing.

### Comparative analyses

This study, as any comparative analysis that considers different species (or other taxonomic units), requires phylogenetically independent comparisons (i.e., comparisons that are evolutionary independent of each other; Felsenstein 1985). Although there is still no phylogeny that includes all finch species, various studies established the relationships within one or a few genera of Fringillidae, including most of its species (Nguembock *et al*. 2009, Zuccon *et al*. 2012, Payevsky 2015). As a result, several genera were shown to be paraphyletic and were divided, causing a reorganization of the taxonomy of the family that is reflected by the Clements *et al*. (2024) checklist that we followed. Current genera are therefore expected to be monophyletic. Based on this rationale, and with the double goal of generating phylogenetically independent comparisons and maximizing the colour elaboration differences between the species of each pair, our family-scale approach consisted in performing single intrageneric comparisons between the species with the most and the least elaborate colour in each genus and assessing whether there is a correlation between plumage and song elaboration. If high-quality recordings were not available for the species with the most or the least elaborate plumage of a genus, the second-most or the second-least elaborate species was selected. When two or more species from a genus had the same CEI and were tied as the most or the least elaborate, the species for which more high-quality recordings were available was selected. This procedure generated 19 intrageneric species pairs that were included in the final dataset for the family-scale analysis (six genera were excluded because their species lacked high-quality recordings).

In addition to these 19 genera with multiple species, 18 genera were represented by a single species (including monospecific genera, genera with only one extant species and genera for which only one species had illustrations in the Handbook of the Birds of the World or pictures in the Macaulay Library). A total of 16 of these genera had high-quality recordings that allowed their potential inclusion in the final dataset for the family-scale analysis. However, in this case we had to form intergeneric species pairs and make sure to include only those species that allowed us to generate pairs that were phylogenetically independent from each other, as well as from the intrageneric species pairs.

To verify the phylogenetic independence of intraspecific pairs, account for any lack of independence, and identify as many phylogenetically independent interspecific pairs as possible, we generated a phylogeny in the birdtree.org website (Jetz *et al*. 2012). This phylogeny was pruned to include 54 species (the 38 species that had the most and least elaborate plumage of their respective genera and the 16 species that are the sole representatives of their genera; Fig. S2). Each species pair (both within genera and among monospecific genera) was considered to be independent from the others if none of the branches of the phylogeny connecting the two species was also used to connect other species pairs (i.e., the species of each pair shared a unique evolutionary history that did not overlap with that of other pairs). This methodology confirmed that our 19 intrageneric species pairs were phylogenetically independent from each other (even though not all genera appear to be monophyletic; see Fig. S2), with the only exception of the genus *Carduelis*. The comparison between the two *Carduelis* species is not independent from that of the two *Spinus* species (i.e., some of the branches that connect *C. corsicana* with *C. carduelis* also connect *S. pinus* with *S. cucullatus*). Because of this, we performed the family-scale analyses both including and excluding *Carduelis*. As expected, removing only one pair of species did not change the outcome of the analyses and we obtained almost identical results with and without *Carduelis*. We therefore decided to retain this genus in the dataset, as it represents one of the most charismatic genera of the family. In addition, we were able to select 8 species from monospecific genera that could form 4 intergeneric pairs independent from each other and from all the intrageneric pairs. Therefore, our final dataset for the family-level analysis had 46 finch species, organized in 19 intrageneric pairs with the species with the most and the least elaborate plumage colour of each genus and 4 intergeneric pairs (Table S5).

In each species pair, we analysed the four aforementioned vocal variables to determine whether the species with the most or the least elaborate plumage invested more energy (song length and syllable production rate), had more complex songs (repertoire index) and exhibited higher vocal performance (vocal deviation). Under the null hypothesis of no association between characters, for each song variable the species with the most elaborate plumage in each pair should have the highest value in 50% of the comparisons and the lowest value in the remaining 50%. A non-parametric binomial test, using the *binom.test* function in the R package *stats* (R Core Team 2023), was performed for each variable to establish if it significantly deviated from these expectations, indicating either a positive or a negative association.

For the genus-scale analyses, we ordered both the *Crithagra* and *Spinus* species in a decreasing CEI order. Species that lacked high-quality recordings were not considered further for the analysis, and as a result the corresponding datasets had 33 *Crithagra* and 17 *Spinus* species. (Tables S6 and S7). We assessed the relationships between the CEI and each acoustic variable using Spearman rank correlations, implemented with the *cor.test* function in R (R Core Team 2024). To account for multiple comparisons, we applied a sequential Bonferroni correction (Holm’s method) using the *p.adjust* function.

## Results

At the family scale (Fringillidae), we did not find a significant association between plumage colour elaboration and any of the four vocalization variables: song length, syllable rate, repertoire index and vocal deviation (Fig. 2; *p* > 0.05 in all cases). In fact, across all variables, in approximately half of the comparisons the species with the most elaborate plumage had more elaborate songs, while in the other half, their songs were simpler.

**Figure 2.**
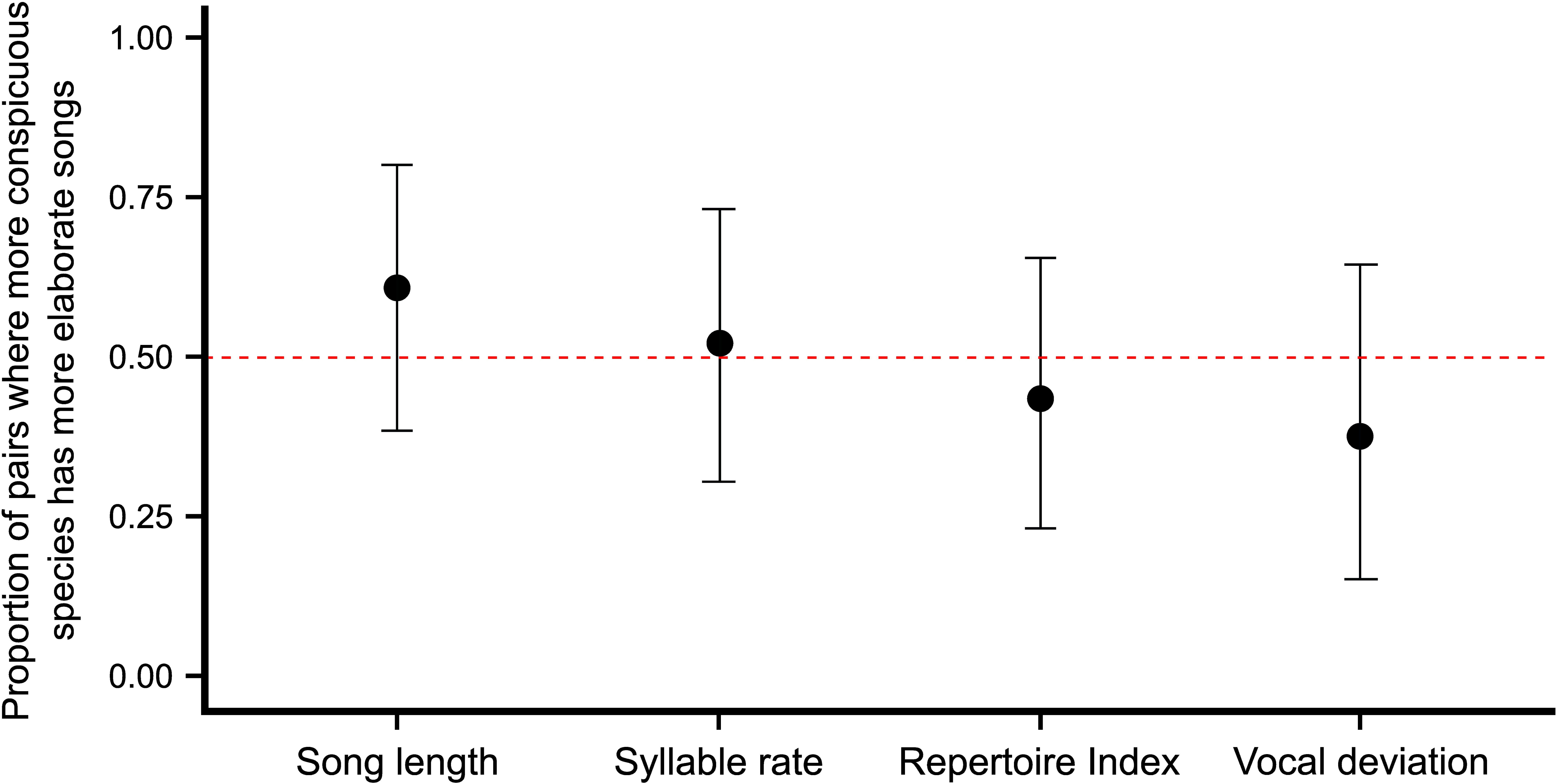
Proportion of species pairs across Fringillidae in which the species with higher CEI exhibits more elaborate songs. Values are plotted with 95% confidence intervals. The dashed red line at 0.50 indicates the expected proportion of pairs in which the species with higher CEI has more elaborate songs if these traits are not correlated. All binomial tests yielded non-significant differences between the observed proportions and those expected by chance (p > 0.05 in all cases).

Consistently, the Spearman rank correlations exhibited a lack of correlation between plumage colour elaboration and the vocal variables at the genus level for both *Crithagra* and *Spinus*. In the case of *Crithagra*, none of the four variables showed an association with the CEI (Fig. 3; Bonferroni-adjusted *p >* 0.05 in all cases). Moreover, the plots show that there are not even tendencies towards either positive or negative correlations. The same is true for *Spinus*, none of the three study variables (as explained in the Methods section, *song length* could not be included for this genus) showed an association or trend (Fig. 4; Bonferroni-adjusted *p* > 0.05 in all cases).

**Figure 3.**
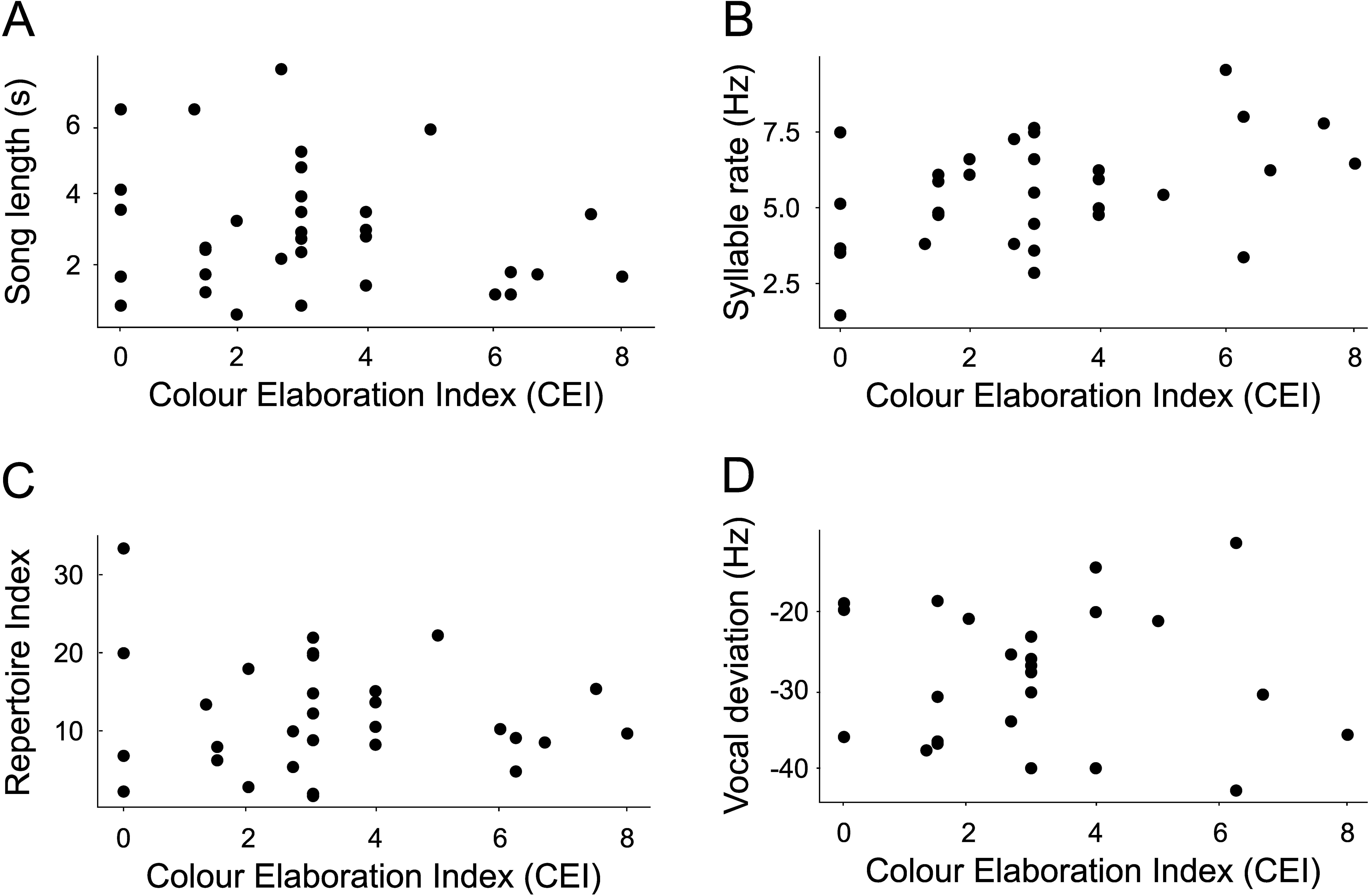
Spearman correlations between the four vocal variables and plumage colour elaboration for the genus *Crithagra*. None of the vocal variables showed a significant correlation with plumage elaboration (*p* > 0.05 in all cases). (A) Song length; (B) Syllable rate; (C) Repertoire index and (D) Vocal deviation.

**Figure 4.**
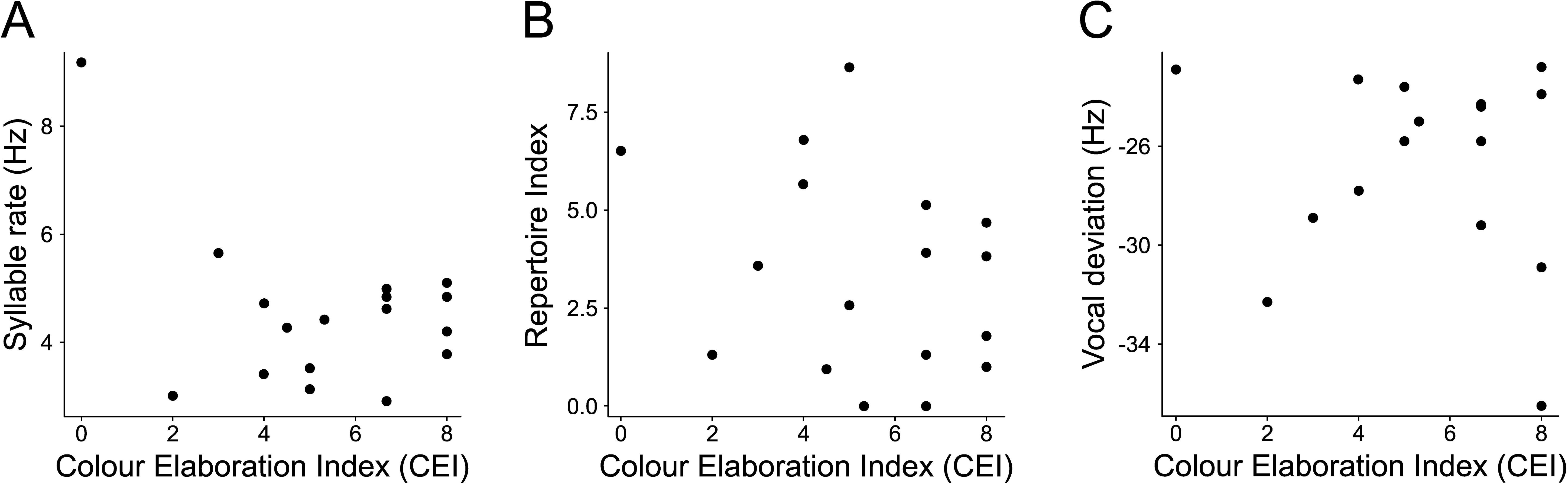
Spearman correlations between the three vocal variables and plumage colour elaboration for the genus *Spinus*. None of the vocal variables showed a significant correlation with colour elaboration (p > 0.05 in all cases). (A) Syllable rate; (B) Repertoire Index and (C) Vocal deviation.

## Discussion

We did not find a correlation between plumage colour elaboration and various aspects of vocal elaboration, including energy investment, song complexity and vocal performance. This holds both for the larger-scale analysis at the family level and for the two assessed genera. Such a result has also been found previously in various avian groups (Ornelas *et al*. 2009, Gonzalez-Voyer 2013, Mason *et al*. 2014, Gomes *et al*. 2017, Matysioková 2017, Marcolin *et al*. 2022), and it is in fact more common than finding a negative (Badyaev *et al*. 2002, Beco *et al*. 2021, Tietze & Hahn 2024) or a positive (Ligon *et al*. 2018, Cárdenas-Posada & Fuxjager 2022, Cardoso *et al*. 2024) correlation between colour and vocal variables. The absence of a correlation between visual and vocal signals has also been documented in other vertebrates, such as tree frogs (Webster *et al*. 2023) and rhesus macaques (Higham *et al*. 2013).

This absence of correlation suggests that colour and vocal elaboration evolve independently in Fringillidae. Contrary to the expectations of the transfer hypothesis (Darwin 1871, Gilliard 1956), this implies that the evolution of an elaborate acoustic or visual signal appears not to affect the complexity of the other signal that is simultaneously expressed by a species. Various non-exclusive reasons could potentially account for this. In the first place, a trade-off in the elaboration of different signals would only be expected if selection operates primarily on single signals, which can replace one another as the target of selection, but this is not necessarily the case (Møller & Pomiankowski 1993, Iwasa & Pomiankowski 1994). In other words, if multiple signals are favoured and they complement each other to indicate the overall quality of the individual, a negative correlation in their elaboration should not be expected. In addition, the transfer hypothesis assumes that both signals are costly. Even though there is clear evidence for the cost of colouring feathers and vocalizing, both in terms of their energetic expenditure (Møller 1996, Oberweger & Goller 2001, Ward *et al*. 2003) and the increased predation risk that they imply (Zuk & Kolluru 1998, Huhta *et al*. 2003, Schmidt & Belinsky 2013, Bliard *et al*. 2020), it is not as clear that more elaborate songs are necessarily more costly (Oberweger & Goller 2001) or that more elaborate plumages always entail higher predation risks (Cain *et al*. 2019).

Both the relevance of expressing multiple signals and the absence of a limiting cost could lead to positive correlations between the elaboration of the signals or to their independent evolution. In particular, if the different signals are assessed by the same receivers (e.g., females) to generate an overall evaluation of the quality of the individual, one would expect sexual selection to operate in the same direction on them, generating a positive correlation between their complexities. This should occur both if the signals complement to provide information about a single quality of the individual (multiple message hypothesis) or if they inform about different aspects of its quality (redundant signal hypotheses) (Møller & Pomiankowski 1993, Johnstone 1996). This appears to be the case in birds of paradise (Ligon *et al*. 2018) and woodpeckers (Cárdenas-Posada & Fuxjager 2022), as well as in other animal groups such as wolf spiders (Hebets *et al*. 2013) and lemurs (Fichtel & Kappeler 2022). To the contrary, if the different signals expressed by the individual are under different selection forces, no correlation should be expected between them. This could be the case if they are associated with different ecological or life-history traits (Gomes *et al*. 2017) or if one of them is targeted by intersexual selection and the other one by intrasexual selection (Shutler & Weatherhead 1990). Even though not finding a significant correlation does not necessarily mean that the association between characters does not exist, the fact that the lack of a correlation has been the most common result in birds strongly suggests that different selection forces could be regularly acting on simultaneously expressed signals.

Other factors could also explain the lack of correlation between the elaboration of different characters, including the fact that large-scale studies might include various lineages with their own characteristics that do present correlations (either negative or positive), but these correlations are lost when the entire group is analysed together (Mason *et al*. 2014). In our study, the lack of an association between colour and vocal elaboration was found both at the family level and in two specific genera (*Crithagra* and *Spinus*), suggesting that this is not an artefact of the large scale of the analysis.

At first glance, our results appear to contradict those of two previous studies of plumage and song association in finches, adding yet more inconsistency to the pattern in this group, as Badyaev *et al*. (2002) found a negative correlation between plumage ornamentation and song complexity in a subgroup of finches and Cardoso *et al*. (2024) found a positive correlation between these traits. However, these different results could be mainly related to differences in the methodology that was employed (Benedict & Najar 2019). Neither plumage nor song elaboration were measured in the same way in these studies. Moreover, phylogenetically independent comparisons were also incorporated in different ways. In fact, this can explain the differences between the results of two of these studies. Cardoso *et al*. (2024) concluded that the positive correlation they observed between colour ornamentation and vocal performance resulted from both traits being associated with body size, rather than from a direct relationship between colour and song. Indeed, once species differences in body size were controlled, colour and song traits were uncorrelated. In this regard, the absence of a correlation in our results mirrors their findings, as our comparisons were largely intrageneric at both taxonomic scales (with only four exceptions involving intergeneric comparisons between monotypic genera), thereby minimizing the effect of size differences among species. In any case, the differences found in these three finch analyses are a clear example of the fact that methodological differences, including the way to determine plumage and song elaboration, could be one of the main reasons for the contrast among studies, irrespective of the bird group assessed in each of them.

In addition to the lack of correlation between plumage and vocal elaboration in finches, our analysis allowed us to confirm that this group produces particularly elaborate songs. This was indicated by their high repertoire indices, the presence of complex syllables in several species and their notable motor performance compared to that of other avian groups. The latter was evidenced by two aspects of finch trills during our measurement of vocal deviation. In the first place, we detected very fast trills in some species, with rates as high as 75 Hz (Fig. S1; note that trills with rates faster than 50 Hz were not included in the vocal deviation analysis, see Methods). This value was even slightly higher than the rates previously detected for Fringillidae (Cardoso *et al*. 2020), in part because of a few species with very fast trills present in our data set that had not been previously analysed in this context (*Fringilla coelebs*, *Mycerobas icterioides*, *Loxia curvirostra*). The second aspect evidencing the notable motor performance of the group is that finches can maintain considerably large bandwidths in their trills as these become faster. Various species had bandwidths as high as 3–5 kHz in trills with rates up to 50 Hz, a much higher value than those of similar trills in other families (Fig. 5), and these large bandwidths were maintained in even faster trills up to 75 Hz (see Fig. S1). This ability of finches to force their phonation system to produce trills that have high syllable rates and large bandwidths suggest that they can relax and contract the syringeal muscles at a higher speed than other families (Laje *et al*. 2002, Mindlin & Laje 2005).

**Figure 5.**
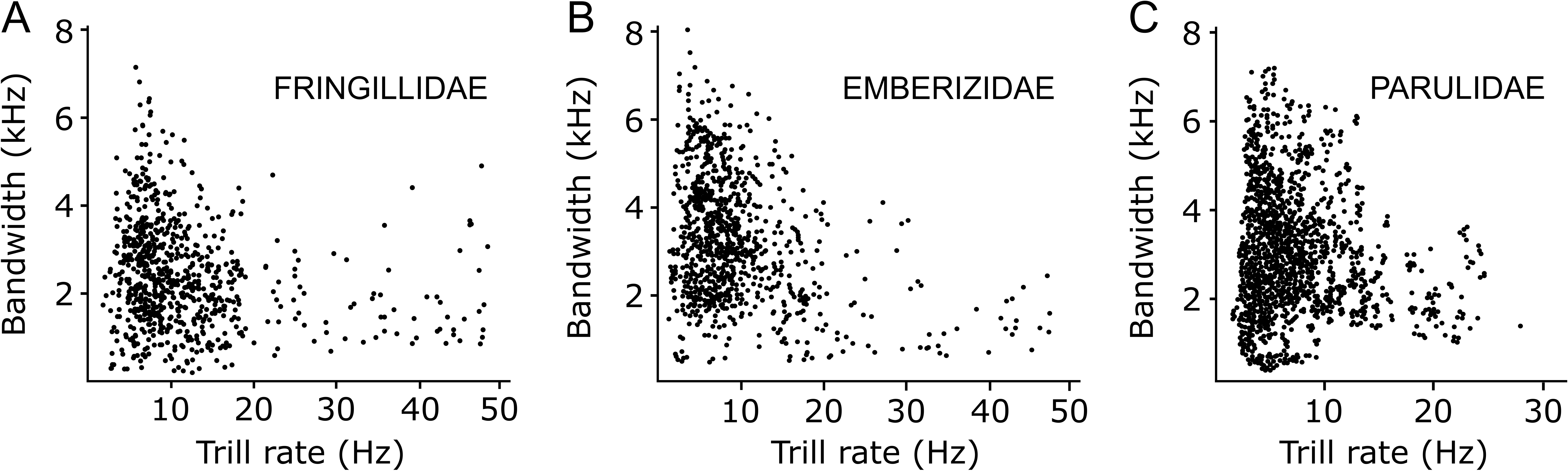
Finches can force their vocal system during trill production compared to other families of birds. The plots show the bandwidths and the syllable rates of the trills included in the songs of various species of three families: (A) Fringillidae, (B) Emberizidae, (C) Parulidae. Finches use considerably larger bandwidths in their fast trills compared with the other two families. Only trills up to 50 Hz were included in plot A. Plots B and C were modified from Podos, 1997 and Cardoso & Hu, 2011, respectively.

This deserves a more detailed study, both in terms of finch anatomy and physiology, as well as the evolutionary processes involved. In the context of the simultaneous use of multiple signals examined in this study, it would be worth investigating if other avian groups also show a lack of correlation between vocal performance and plumage colour elaboration (or other elaborate traits). This would allow assessing whether the absence of a trade-off between these signals in finches is related to their apparent ability to force their phonation system, possibly incurring a lower associated cost than other birds (but see Gomes *et al*. 2017, for an example of lack of association between vocal performance and colouration in estrildid finches). Finally, it is worth mentioning that we found a similar relation between bandwidth and trill rate across a broad range of trill speeds that could potentially include both mini-breaths and pulsatile expiration, as well as a different cycling patterns of bronchial air pressure and syrinx muscle tension (Laje *et al*. 2002, Mindlin & Laje 2005, Lijtmaer 2008). Analyses of finch respiratory patterns, bronchial air pressures and muscle tension during song production at varying syllable rates could clarify this aspect of their communication.

## Conclusions

In sum, our results taken together suggest that finch plumage and vocal elaboration evolved independently, in spite of the fact that they produce highly elaborate songs. This study thus adds to the growing list of analyses indicating that, although contrasting results have been obtained, plumage and vocal elaboration tend to evolve independently in birds. Disparate results have also been found in other animal groups (e.g., Hebets *et al*. 2013, Higham *et al*. 2013, Fichtel & Kappeler 2022, Webster *et al*. 2023). This highlights the complexity of multimodal signaling and emphasizes that broad comparative analyses of simultaneously expressed traits across different taxonomic groups are needed to advance in the understanding of the evolution of secondary sexual characters and animal communication.

## Acknowledgments

We thank the members of the Ornithology Division of the Museo Argentino de Ciencias Naturales ‘Bernardino Rivadavia’ and Gabriel Mindlin for their helpful discussions that enriched various aspects of this study. This research was supported by the Consejo Nacional de Investigaciones Científicas y Técnicas (CONICET, Argentina), the Agencia Nacional de Promoción de la Investigación, el Desarrollo Tecnológico y la Innovación (Agencia I+D+i, Argentina) and the Richard Lounsbery Foundation. We also thank the Macaulay Library and Xeno-Canto for providing access to the bird song recordings analyzed in this study. We are grateful to Patricio I. Casale and Jazmín Grillo Balboa for their assistance in the design of the figures.

