## Supporting Information for "Evaluating the association between vocalizations and colouration in finches (Fringillidae), a passerine group with elaborate signals"

### Table of Contents:


| | |
|---|---|
| **Figure S1** | Page 2 |
| **Figure S2** | Page 3 |
| **Table S1** | Page 5 |
| **Table S2** | Page 7 |
| **Table S3** | Page 9 |
| **Table S4** | Page 11 |
| **Table S5** | Page 12 |
| **Table S6** | Page 14 |
| **Table S7** | Page 16 |

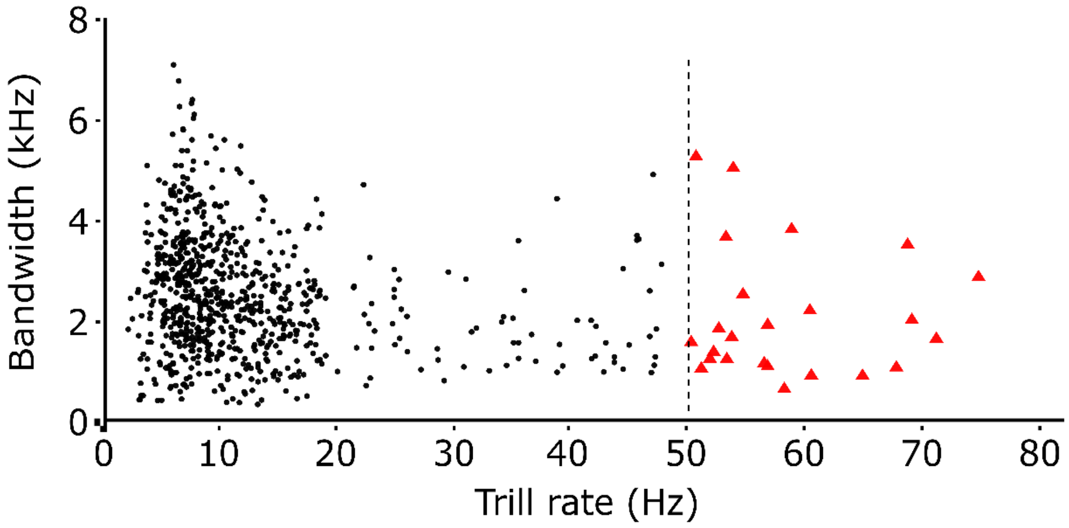


**Figure S1**. Finches are capable of forcing their vocal system during their trills. The plot shows that they can produce trills with rates up to 75 Hz and that they can use large bandwidths (up to 5 kHz) in these fast trills. Black dots: trills with rates up to 50 Hz that were used in the vocal deviation analysis; Red triangles: trills with rates faster than 50 Hz that were not included in the vocal deviation analysis because their production mechanism is thought to be different and less constrained.

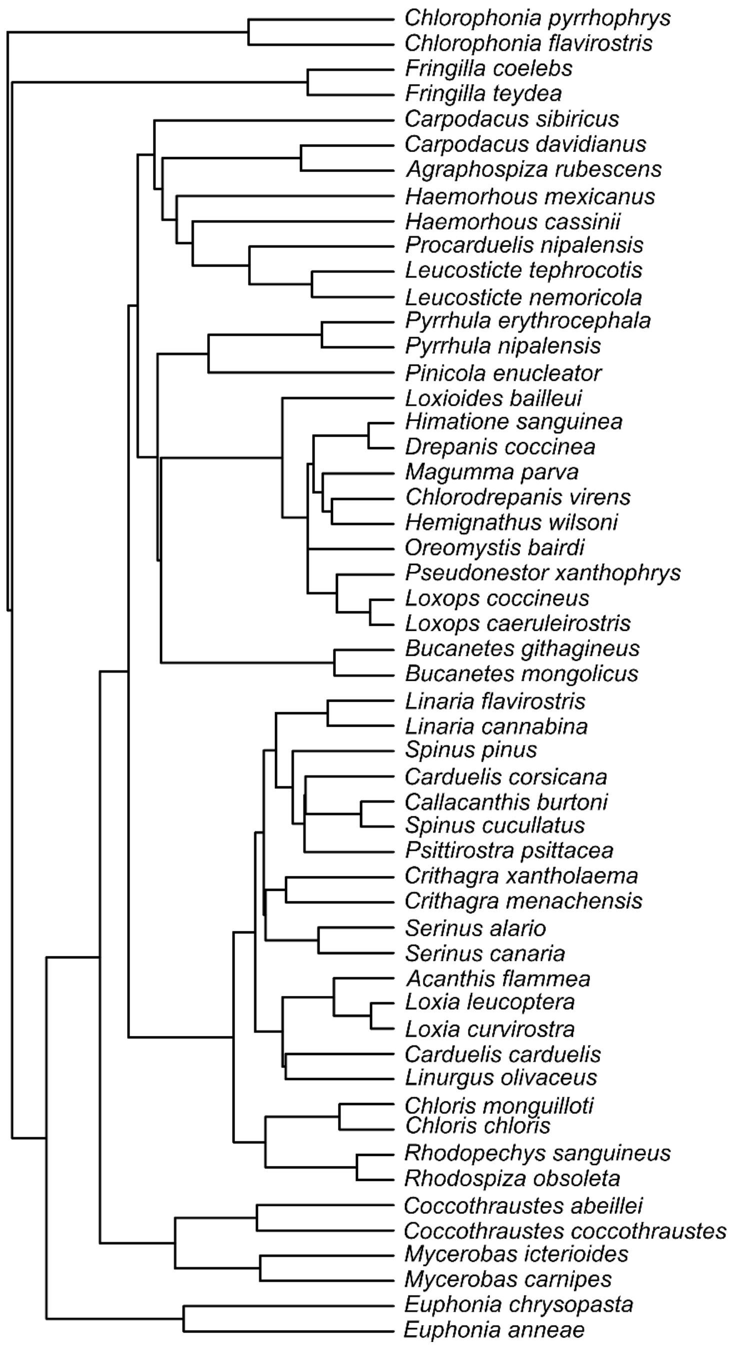


**Figure S2.** Phylogeny obtained from the birdtree.org website (Jetz et al., 2012) pruned to include only the species used for the family-level analysis of Fringillidae. The topology of this phylogeny was used to confirm that the pairs of species compared were independent from each other. Note that the only exception is the comparison between the two *Carduelis* species, which is not independent from the two *Spinus* comparisons (i.e., some of the branches that connect *C. corsicana* with *C. carduelis* also connect *S. pinus* with *S. cucullatus*). This is why we repeated the analysis without the genus *Carduelis*, obtaining the same results as with the complete dataset.

**Table S1**. Fringillidae species for which plumage conspicuousness was analysed (i.e., all extant species that had either illustrations in the Handbook of the Birds of the World (Collar et al., 2010)) or pictures in the Macaulay Library (http://macaulaylibrary.org/)) and their Colour Elaboration Index (CEI) value. Species highlighted in bold were used for the family-level analysis (see the Methods section for details about the selection of species for comparison).

| Genus | CEI |
|---|---|
| ***Fringilla*** | ***F. coelebs*: 14.00**; *F. maderensis*: 10.50; *F. spodiogenys*: 9.03; *F. polatzeki*: 2.00; *F. moreletti*: 10.80; *F. canariensis*: 10.00; ***F. teydea*: 2.00**; *F. montifringilla*: 10.00 |
| ***Chlorophonia*** | *C. elegantissima*: 10.00; *C. musica*: 10.00; *C. sclateri*: 10.00; *C. flavifrons*: 6.25; *C. cyanocephala*: 8.00; *C. cyanea*: 12.00; ***C. pyrrhophrys*: 18.00**; ***C. flavirostris*: 8.00**; *C. occipitalis*: 10.00; *C. callophrys*: 14.00 |
| ***Euphonia*** | *E. jamaica*: 2.00; *E. saturata*: 6.00; *E. plumbea*: 2.00; *E. chlorotica*: 6.00; *E. finschi*: 6.00; *E. concinna*: 6.00; *E. trinitatis*: 6.00; *E. godmani*: 6.00; *E. affinis*: 6.00; *E. luteicapilla*: 6.00; ***E. chrysopasta*: 1.00**; *E. minuta*: 8.00; *E. chalybea*: 6.00; *E. violacea*: 6.00; *E. hirundinacea*: 6.00; *E. laniirostris*: 6.00; *E. gouldi*: 5.32; *E. fulvicrissa*: 6.00; ***E. anneae*: 8.00**; *E. xanthogaster*: 6.00; *E. mesochrysa*: 3.00; *E. cayennensis*: 4.00; *E. rufiventris*: 4.00; *E. pectoralis*: 6.00; *E. imitans*: 6.00. |
| ***Mycerobas*** | ***M. icterioides*: 6.00**; *M. affinis*: 6.00; *M. melanozanthos*: 6.00; ***M. carnipes*: 8.00**. |
| ***Coccothraustes*** | ***C. abeillei*: 6.68**; *C. vespertinus*: 10.00; ***C. coccothraustes*: 20.00**. |
| ***Chlorodrepanis*** | ***C. virens*: 1.00**; ***C. flava*: 2.00**; *C. stejnegeri*: 1.00. |
| ***Loxops*** | *L. mana*: 1.00; ***L. caeruleirostris*: 1.00**; ***L. coccineus*: 2.00**; *L. wolstenholmei*: 2.00; *L. ochraceus*: 2.00. |
| ***Carpodacus*** | *C. erythrinus*: 1.50; *C. sipahi*: 4.00; *C. ferreorostris*: 1.00; *C. rhodochlamys*: 0.50; *C. grandis*: 1.00; *C. pulcherrimus*: 2.00; ***C. davidianus*: 1.00**; *C. waltoni*: 2.00; *C. edwardsii*: 2.00; *C. rodochroa*: 2.00; *C. rodopeplus*: 3.00; *C. verreauxii*: 3.00; *C. vinaceus*: 4.50; *C. formosanus*: 4.50; *C. synoicus*: 3.00; *C. stoliczkae*: 4.50; *C. roborowskii*: 1.50; *C. sillemi*: 1.50; *C. rubicilloides*: 2.00; *C. rubicilla*: 2.00; ***C. sibiricus*: 4.50**; *C. puniceus*: 4.50; *C. subhimachalus*: 3.00; *C. roseus*: 3.00; *C. trifasciatus*: 7.50; *C. thura*: 4.00; *C. dubius*: 4.50. |
| ***Pyrrhula*** | ***P. nipalensis*: 4.00**; *P. leucogenis*: 8.00; *P. aurantiaca*: 6.68; ***P. erythrocephala*: 10.00**; *P. erythaca*: 10.00; *P. owstoni*: 6.00; *P. murina*: 6.00; *P. pyrrhula*: 8.00. |
| ***Bucanetes*** | ***B. githagineus*: 0.50**; ***B. mongolicus*: 3.00**. |
| ***Leucosticte*** | ***L. nemoricola*: 1.00**; *L. brandti*: 2.00; *L. arctoa*: 1.00; ***L. tephrocotis*: 8.75**; *L. atrata*: 6.68; *L. australis*: 4.00. |
| ***Haemorhous*** | ***H. mexicanus*: 4.50**; *H. purpureus*: 4.00; ***H. cassinii*: 4.00**. |
| ***Chloris*** | ***C. chloris*: 2.00**; *C. sinica*: 6.25; *C. spinoides*: 7.50; ***C. monguilloti*: 7.50**; *C. ambigua*: 6.00. |
| ***Crithagra*** | *C. leucopygia*: 1.00; *C. mozambica*: 6.68; *C. citrinelloides*: 4.00; *C. frontalis*: 4.00; *C. hyposticta*: 3.00; *C. capistrata*: 4.00; *C. koliensis*: 1.50; *C. scotops*: 4.00; *C. rothschildi*: 0.50; *C. atrogularis*: 3.00; *C. reichenowi*: 3.00; *C. xanthopygia*: 2.00; *C. citrinipectus*: 7.50; *C. dorsostriata*: 6.25; *C. flavigula*: 6.00; ***C. xantholaema*: 8.00**; *C. donaldsoni*: 3.00; *C. buchanani*: 1.50; *C. sulphurata*: 4.00; |

| | |
|---|---|
| | *C. flaviventris*: 5.00; *C. albogularis*: 3.00; *C. striolata*: 1.50; *C. whytii*: 6.68; *C. burtoni*: 3.00; *C. melanochroa*: 1.50; *C. rufobrunnea*: 0.50; *C. concolor*: 0.50; *C. leucoptera*: 2.68; *C. mennelli*: 6.25; *C. canicapilla*: 2.68; *C. gularis*: 3.00; *C. striatipectus*: 1.32; *C. reichardi*: 1.32; *C. tristriata*: 2.00; ***C. menachensis*: 0.50**; *C. ankoberensis*: 1.00; *C. totta*: 3.00; *C. symonsi*: 1.50. |
| ***Linaria*** | ***L. flavirostris*: 0.50**; ***L. cannabina*: 5.00**; *L. yemenensis*: 5.00; *L. johannis*: 4.00. |
| ***Loxia*** | *L. pytyopsittacus*: 2.00; *L. scotica*: 2.00; ***L. curvirostra*: 2.00**; *L. sinesciuris*: 2.00; *L. megaplaga*: 4.50; ***L. leucoptera*: 6.00**. |
| ***Carduelis*** | ***C. carduelis*: 14.88**; *C. caniceps*: 10.00; *C. citrinella*: 5.32; ***C. corsicana*: 5.32**. |
| ***Serinus*** | *S. serinus*: 2.00; ***S. canaria*: 2.00**; *S. pusillus*: 6.00; *S. syriacus*: 2.00; *S. flavivertex*: 2.00; *S. canicollis*: 2.00; *S. nigriceps*: 8.00; ***S. alario*: 8.00**. |
| ***Spinus*** | *S. thibetanus*: 2.00; *S. spinus*: 5.00; ***S. pinus*: 1.5**; *S. atriceps*: 5.32; *S. notatus*: 8; *S. dominicensis*: 6.00; *S. psaltria*: 3.00; *S. lawrencei*: 8.00; *S. tristis*: 5.00; *S. spinescens*: 6.68; *S. yarrellii*: 8.00; *S. xanthogastrus*: 3.99; ***S. cucullatus*: 8.00**; *S. crassirostris*: 6.68; *S. magellanicus*: 6.68; *S. siemiradzkii*: 8.00; *S. olivaceus*: 8.00; *S. atratus*: 4.00; *S. uropygialis*: 4.50; *S. barbatus*: 6.68. |
| ***Rhodopechys*** | ***R. sanguineus*: 3.00** |
| ***Loxioides*** | ***L. bailleui*: 4.50** |
| ***Pinicola*** | ***P. enucleator*: 5.32** |
| ***Rhodospiza*** | ***R. obsoleta*: 2.00** |
| ***Oreomystis*** | ***O. bairdi*: 1.00** |
| ***Pseudonestor*** | ***P. xanthophrys*: 4.50** |
| ***Himatione*** | ***H. sanguinea*: 4.00** |
| ***Drepanis*** | ***D. coccinea*: 6.00** |

**Table S2**. Number of individuals of each species used for the Fringillidae-family scale analysis and their corresponding Xeno-Canto and Macaulay Library recording catalogue numbers.

| Species | N | Catalogue Number |
|---|---|---|
| *Fringilla coelebs* | 10 | ML187165, ML36192, ML86162, ML241035, ML187143, ML141478, ML248519, ML202479, ML246607, ML187154 |
| *Fringilla teydea* | 3 | ML20579, XC131868, XC349387 |
| *Carpodacus sibiricus* | 10 | ML142905421, ML203970901, ML389157101, ML389157111, XC61925, XC183349, XC505257, XC832414, XC916118, XC1006257 |
| *Carpodacus davidianus* | 2 | ML251379271, ML457889631 |
| *Carduelis carduelis* | 8 | ML12751, ML86201, ML104977, ML162878, ML240798, ML248454, ML248535, ML248681 |
| *Carduelis corsicana* | 8 | XC134339, XC146345, XC349878, XC448044, XC470702, XC471879, XC471925, XC471929 |
| *Chlorophonia pyrrhophris* | 4 | ML238215, ML269865, XC87041, XC102288 |
| *Chlorophonia flavirostris* | 3 | XC13315, XC261143, ML262532 |
| *Euphonia anneae* | 4 | ML134152061, ML633923872, XC75588, XC538981 |
| *Euphonia chrysopasta* | 10 | ML34249, ML48672, ML70752, ML74759, ML88282, ML94901, ML244119, ML245150, ML245342, ML261038 |
| *Mycerobas icterioides* | 8 | ML24754, ML41930, ML179417, ML179442, ML179444, XC19577, XC255922, XC311728 |
| *Mycerobas carnipes* | 2 | XC19920, XC35566 |
| *Coccothraustes coccothraustes* | 6 | XC112670, XC363825, XC403252, XC468472, XC411225, XC213839 |
| *Coccothraustes abeillei* | 7 | ML12953, ML86675, ML145972, ML145975, ML145984, ML145989, ML215805 |
| *Chlorodrepanis virens* | 10 | ML5270, ML5277, ML5283, ML5838, ML5842, ML6071, ML218676, ML236859, ML234845, ML218684 |
| *Chlorodrepanis flava* | 6 | XC59050, XC59051, XC59052, XC174828, XC174829, XC454888 |
| *Loxops coccineus* | 10 | ML5257, ML5891, ML129226, ML129229, ML129259, ML129260, ML129262, ML129265, ML5876, ML6046 |
| *Loxops caerulirostris* | 8 | XC27306, XC27320, XC174954, XC210201, ML56464, ML234885, ML234895, ML234892 |
| *Pyrrhula erythrocephala* | 8 | ML41586, ML41895, ML53568, XC119995, XC256464, XC319293, XC119994, XC118249 |
| *Pyrrhula nipalensis* | 9 | ML12997, ML12998, ML68741, ML73616, ML140818, ML175049, ML236443, XC35451, XC398310 |
| *Bucanetes githagineus* | 4 | XC164199, XC354891, XC358584, XC490889 |
| *Bucanetes mongolicus* | 2 | XC133401, ML197553 |
| *Leucosticte nemoricola* | 5 | XC108741, XC145255, XC426653, XC426654, XC491299 |
| *Leucosticte tephrocotis* | 5 | XC91905, XC143944, XC143945, XC143947, XC352924 |

| | | |
|---|---|---|
| *Haemorhous mexicanus* | 10 | ML27129, ML27141, ML27159, ML27161, ML27164, ML27172, ML27176, ML27180, ML27871, ML539584 |
| *Haemorhous cassinii* | 8 | ML80302, ML80328, ML90329, ML80363, ML87913, ML96371, ML96372, ML99333 |
| *Chloris chloris* | 9 | XC478729, XC480880, XC486002, XC486183, XC489263, XC493882, XC495535, ML36480, ML204481 |
| *Chloris monguilloti* | 5 | XC26288, XC26534, XC126964, XC126968, XC201082 |
| *Crithagra xanthoalema* | 4 | ML100211, ML28073981, XC210309, XC210310 |
| *Crithagra menachensis* | 3 | XC44563, XC44564, XC71576 |
| *Linaria cannabina* | 7 | XC467508, XC467969, XC468810, XC477710, XC481813, XC483247, XC500408 |
| *Linaria flavirostris* | 5 | XC100476, XC189577, XC246297, XC364664, XC414433 |
| *Loxia curvirostra* | 10 | ML34743, ML86240, ML137498, ML138303, ML138303, ML138308, ML138309, ML138310, ML138318, ML233285 |
| *Loxia leucoptera* | 8 | ML27037, ML42389, ML112164, ML112170, ML112172, ML133395, ML516586, ML516594 |
| *Serinus canaria* | 8 | XC79800, XC349868, XC354850, XC366507, XC367178, XC374955, XC374956, XC398574 |
| *Serinus alario* | 5 | XC391059, XC392692, XC392694, XC392696, XC392697 |
| *Spinus pinus* | 4 | ML516482, ML217604, ML205799, ML99395 |
| *Spinus cucullatus* | 2 | ML111830, ML111840 |
| *Pseudonestor xanthophrys* | 3 | ML236862, ML234846, ML6077 |
| *Oreomystis bairdi* | 1 | ML90524951 |
| *Pinicola enucleator* | 4 | ML516416, ML516465, ML516522, ML516579 |
| *Loxioides bailleui* | 7 | ML5195, ML32525, ML129232, ML129234, ML183507, ML234875, ML234877 |
| *Drepanis coccinea* | 3 | ML94200351, ML5859, ML129127 |
| *Himatione sanguinea* | 8 | ML10932, ML129118, ML129147, ML234848, ML632682749, ML636917596, ML632487435, ML632649403 |
| *Rhodopechys sanguineus* | 4 | XC141509, XC350469, ML104940, ML220029 |
| *Rhodospiza obsoleta* | 5 | XC134470, XC306834, XC471582, XC471587, XC471597 |

**Table S3**. Number of individuals of each species used for the analysis of the genus *Crithagra* and their corresponding Xeno-Canto and Macaulay Library recording catalogue numbers.

| **Species** | **N** | **Catalogue Number** |
|---|---|---|
| *Crithagra menachensis* | 3 | XC44563, XC44564, XC71756 |
| *Crithagra rothschildi* | 3 | ML203934381, XC298737, XC447318 |
| *Crithagra rufobrunnea* | 3 | ML96203, ML102119, XC339870 |
| *Crithagra concolor* | 1 | XC17889 |
| *Crithagra leucopygia* | 1 | XC317109 |
| *Crithagra reichardi* | 3 | ML277084, ML277142, ML277144 |
| *Crithagra koliensis* | 1 | XC538098 |
| *Crithagra buchanani* | 4 | ML280267, ML280270, ML280272, XC397519 |
| *Crithagra striolata* | 10 | ML1560, ML13021, ML17945, ML17972, ML17974, ML276139, ML276477, ML276579, ML276580, ML33720801 |
| *Crithagra symonsi* | 3 | ML275697, ML275710, ML176711871 |
| *Crithagra xanthopygia* | 4 | ML276989, ML31283581, XC209972, XC300224 |
| *Crithagra tristriata* | 3 | ML276486, ML190842571, XC268727 |
| *Crithagra leucoptera* | 9 | ML278435, ML278436, ML123444471, ML177674051, ML177674111, ML187468311, XC392454, XC392455, XC395388 |
| *Crithagra canicapilla* | 1 | ML13013 |
| *Crithagra hyposticta* | 3 | ML49648, XC396039, XC510114 |
| *Crithagra atrogularis* | 3 | ML123443831, XC324307, XC498688 |
| *Crithagra reichenowi* | 10 | ML58518, ML58521, ML276622, ML276623, ML276649, ML276805, ML280586, XC83676, XC300215, XC300222 |
| *Crithagra donaldsoni* | 7 | ML276735, ML276738, ML276745, XC82653, XC82654, XC267119, XC306594 |
| *Crithagra albogularis* | 10 | ML277604, ML278274, ML278300, ML278301, ML278376, XC346755, XC392436, XC392438, XC392440, XC459452 |
| *Crithagra burtoni* | 2 | ML1279, ML135410291 |
| *Crithagra gularis* | 10 | ML278243, ML278244, ML278554, ML278555, ML278559, ML110836271, XC392452, XC496887, XC521817, XC523725 |
| *Crithagra totta* | 2 | ML212083001, XC523740 |
| *Crithagra citrinelloides* | 1 | XC294336 |
| *Crithagra frontalis* | 6 | ML1242, ML1544, ML1545, ML1559, XC93584, XC290019 |
| *Crithagra scotops* | 2 | ML275777, XC492730 |
| *Crithagra sulphurata* | 4 | ML49630, ML278647, XC62663, XC512680 |
| *Crithagra flaviventris* | 3 | ML278305, ML278595, XC451936 |
| *Crithagra flavigula* | 7 | ML276228, ML276223, ML31280651, XC300233, XC307073, XC445059, XC266664 |
| *Crithagra dorsostriata* | 4 | XC268719, XC268721, XC307043, XC450583 |

| | | |
|---|---|---|
| *Crithagra mennelli* | 2 | ML278837, ML278838 |
| *Crithagra mozambica* | 10 | ML275397, ML275944, ML275949, ML277270, ML279331, ML279332, ML279333, ML280574, ML280669, ML206673441 |
| *Crithagra citrinipectus* | 2 | XC395991, XC395992 |
| *Crithagra xantholaema* | 4 | ML100211, ML28073981, XC210309, XC210310 |

**Table S4**. Number of individuals of each species used for the analysis of the genus *Spinus* and their corresponding Xeno-Canto and Macaulay Library recording catalogue numbers.

| Species | N | Catalogue Number |
|---|---|---|
| *Spinus pinus* | 9 | ML440873361, ML433059221, ML347615341, ML324129201, ML314453951, ML199464001, ML153777441, ML149060331, ML148712721 |
| *Spinus thibetanus* | 1 | ML223892221 |
| *Spinus psaltria* | 10 | ML533959041, ML490950751, ML465336241, ML394908051, ML394908021, ML368196361, ML359464871, ML213644261, ML149096321, ML79049701 |
| *Spinus xanthogastrus* | 3 | ML230866161, ML165874, ML63566 |
| *Spinus atratus* | 6 | ML520930331, ML301165, ML204006781, ML515484, ML210219, ML146471 |
| *Spinus uropygialis* | 1 | ML440880051 |
| *Spinus spinus* | 5 | ML461957921, ML425126931, ML305035, ML65424791, ML51992321 |
| *Spinus tristis* | 10 | ML531247211, ML527841051, ML502733961, ML472492451, ML466334221, ML463572751, ML458963571, ML454222151, ML448455931, ML442476451 |
| *Spinus atriceps* | 1 | ML217539 |
| *Spinus spinescens* | 3 | ML351676691, ML286104, ML63546 |
| *Spinus crassirostris* | 5 | ML203960101, ML238376, ML240377, ML171464, ML146474 |
| *Spinus magellanicus* | 10 | ML514648701, XC47334, ML278031461, ML67956091, ML57032621, ML523236, ML515454, ML514684, ML197291, ML174877 |
| *Spinus barbatus* | 6 | ML303623, ML202226081, ML140682391, ML118512151, ML68834481, ML146627 |
| *Spinus notatus* | 3 | ML147689111, ML104332271, ML164101 |
| *Spinus lawrencei* | 3 | ML487910991, ML457331991, ML212095771 |
| *Spinus olivaceus* | 2 | ML35819, ML17780 |
| *Spinus cucullatus* | 2 | ML111830, ML111840 |

**Table S5**. Colour Elaboration Index and vocal trait values for each species included in the family-level analysis of Fringillidae. CEI: Colour Elaboration Index; RI: Repertoire Index. Song length is expressed in seconds and syllable rate and vocal deviation are expressed in Hz. Species with dashes in the "Vocal deviation" column indicate songs that did not contain trills.

| **Species** | **CEI** | **Song length** | **Syllable rate** | **RI** | **Vocal deviation** |
|---|---|---|---|---|---|
| *Fringilla coelebs* | 14.00 | 2.49 | 9.32 | 3.94 | -23.61 |
| *Fringilla teydea* | 2.00 | 1.96 | 4.99 | 4.33 | -49.17 |
| *Carpodacus sibiricus* | 4.50 | 1.13 | 7.75 | 6.51 | - |
| *Carpodacus davidianus* | 1.00 | 0.67 | 7.73 | 2.23 | - |
| *Carduelis carduelis* | 14.88 | 2.40 | 10.08 | 6.21 | -38.61 |
| *Carduelis corsicana* | 5.32 | 0.65 | 8.57 | 1.63 | -63.35 |
| *Chlorophonia pyrrhophris* | 18.00 | 1.19 | 6.49 | 4.92 | - |
| *Chlorophonia flavirostris* | 8.00 | 0.57 | 1.85 | 1.00 | - |
| *Euphonia anneae* | 8.00 | 1.12 | 4.74 | 1.47 | -45.77 |
| *Euphonia chrysopasta* | 0.00 | 0.71 | 9.38 | 5.30 | -27.88 |
| *Mycerobas carnipes* | 8.00 | 0.47 | 8.28 | 2.00 | -47.02 |
| *Mycerobas icterioides* | 6.00 | 0.61 | 9.24 | 2.88 | -51.09 |
| *Coccothraustes coccothraustes* | 20.00 | 3.88 | 2.02 | 4.00 | -61.03 |
| *Coccothraustes abeillei* | 6.68 | 0.96 | 8.95 | 2.52 | -49.17 |
| *Chlorodrepanis flava* | 2.00 | 1.72 | 6.04 | 1.00 | -47.25 |
| *Chlorodrepanis virens* | 1.00 | 2.09 | 5.07 | 1.10 | -22.97 |
| *Loxops caerulirostris* | 1.00 | 0.51 | 5.32 | 1.13 | -22.97 |
| *Loxops coccineus* | 2.00 | 1.85 | 6.68 | 1.90 | -57.80 |
| *Pyrrhula erythrocephala* | 10.00 | 0.39 | 5.42 | 2.00 | - |
| *Pyrrhula nipalensis* | 4.00 | 0.88 | 7.72 | 2.56 | - |
| *Bucanetes mongolicus* | 3.00 | 0.78 | 2.64 | 2.00 | - |
| *Bucanetes githagineus* | 0.50 | 0.62 | 4.99 | 1.00 | - |
| *Leucosticte tephrocotis* | 8.75 | 0.30 | 8.06 | 1.50 | - |
| *Leucosticte nemoricola* | 1.00 | 0.32 | 9.56 | 2.00 | - |
| *Haemorhous mexicanus* | 4.50 | 2.52 | 6.21 | 12.25 | -55.51 |
| *Haemorhous cassinii* | 4.00 | 2.57 | 7.04 | 14.44 | -45.13 |
| *Chloris monguilloti* | 7.50 | 0.45 | 12.6 | 2.61 | -53.09 |
| *Chloris chloris* | 2.00 | 1.64 | 12.1 | 1.93 | -42.30 |
| *Crithagra xantholaema* | 8.00 | 1.69 | 6.31 | 0.70 | -53.21 |
| *Crithagra menachensis* | 0.50 | 0.76 | 3.68 | 1.92 | -50.65 |

| | | | | | |
|---|---|---|---|---|---|
| *Linaria cannabina* | 5.00 | 2.85 | 11.01 | 7.90 | -38.18 |
| *Linaria flavirostris* | 0.50 | 1.82 | 6.98 | 5.70 | -24.93 |
| *Loxia leucoptera* | 6.00 | 4.46 | 11.05 | 3.94 | -39.19 |
| *Loxia curvirostra* | 2.00 | 2.36 | 6.03 | 3.26 | -48.74 |
| *Serinus alario* | 8.00 | 2.13 | 9.89 | 4.67 | -51.13 |
| *Serinus canaria* | 2.00 | 3.41 | 5.52 | 3.84 | -42.80 |
| *Spinus cucullatus* | 8.00 | 1.36 | 9.18 | 1.00 | -45.90 |
| *Spinus pinus* | 1.50 | 2.00 | 5.65 | 4.15 | -39.74 |
| *Pseudonestor xanthophrys* | 4.50 | 1.82 | 4.41 | 3.26 | -36.17 |
| *Oreomystis bairdi* | 1.00 | 1.55 | 5.36 | 1.00 | -44.04 |
| *Pinicola enucleator* | 5.32 | 0.89 | 7.60 | 2.68 | - |
| *Loxioides bailleui* | 4.50 | 0.55 | 8.92 | 3.57 | - |
| *Drepanis coccinea* | 6.00 | 2.76 | 3.09 | 3.32 | - |
| *Himatione sanguinea* | 4.00 | 1.99 | 6.02 | 6.00 | - |
| *Rhodopechys sanguineus* | 3.00 | 0.92 | 7.78 | 4.55 | -57.31 |
| *Rhodospiza obsoleta* | 2.00 | 1.37 | 6.84 | 5.00 | -29.22 |

**Table S6.** Colour Elaboration Index and vocal trait values for each species included in the genus-level analysis of *Crithagra*. CEI: Colour Elaboration Index; RI: Repertoire Index. Song length is expressed in seconds and syllable rate and vocal deviation are expressed in Hz. Species with dashes in all their vocal traits could not be analysed due to the lack of high-quality recordings. Species with dashes only in the "Vocal deviation" column indicate songs that did not contain trills.

| **Species** | **CEI** | **Song length** | **Syllable rate** | **RI** | **Vocal deviation** |
|---|---|---|---|---|---|
| *Crithagra menachensis* | 0.50 | 0.78 | 3.52 | 1.92 | -36.40 |
| *Crithagra rothschildi* | 0.50 | 4.06 | 7.34 | 19.40 | -20.32 |
| *Crithagra rufobrunnea* | 0.50 | 3.46 | 3.36 | 6.42 | -19.51 |
| *Crithagra concolor* | 0.50 | 1.58 | 1.28 | 2.00 | - |
| *Crithagra leucopygia* | 1.00 | 6.29 | 4.99 | 32.80 | - |
| *Crithagra ankoberensis* | 1.00 | - | - | - | - |
| *Crithagra striatipectus* | 1.32 | - | - | - | - |
| *Crithagra reichardi* | 1.32 | 6.26 | 3.64 | 12.84 | -38.12 |
| *Crithagra koliensis* | 1.50 | 1.14 | 5.72 | 5.95 | -37.07 |
| *Crithagra buchanani* | 1.50 | 2.36 | 4.68 | 7.55 | -37.31 |
| *Crithagra striolata* | 1.50 | 2.42 | 5.94 | 7.70 | -31.40 |
| *Crithagra melanochroa* | 1.50 | - | - | - | - |
| *Crithagra symonsi* | 1.50 | 1.62 | 4.59 | 5.75 | -19.26 |
| *Crithagra xanthopygia* | 2.00 | 3.15 | 6.44 | 17.40 | -21.54 |
| *Crithagra tristriata* | 2.00 | 0.51 | 5.97 | 2.43 | - |
| *Crithagra leucoptera* | 2.68 | 7.44 | 3.68 | 4.92 | -26.10 |
| *Crithagra canicapilla* | 2.68 | 2.08 | 7.12 | 9.60 | -34.40 |
| *Crithagra hyposticta* | 3.00 | 3.81 | 4.33 | 19.24 | - |
| *Crithagra atrogularis* | 3.00 | 5.10 | 5.39 | 21.56 | -28.19 |
| *Crithagra reichenowi* | 3.00 | 2.65 | 5.37 | 14.23 | -23.83 |
| *Crithagra donaldsoni* | 3.00 | 3.39 | 3.47 | 1.71 | -40.41 |
| *Crithagra albogularis* | 3.00 | 4.64 | 7.34 | 19.47 | -30.80 |
| *Crithagra burtoni* | 3.00 | 0.77 | 2.67 | 1.50 | -26.60 |
| *Crithagra gularis* | 3.00 | 2.85 | 6.44 | 8.41 | -27.47 |
| *Crithagra totta* | 3.00 | 2.25 | 7.53 | 11.80 | -23.77 |
| *Crithagra citrinelloides* | 4.00 | 1.33 | 5.81 | 7.81 | - |
| *Crithagra frontalis* | 4.00 | 2.69 | 4.63 | 13.32 | -15.07 |
| *Crithagra capistrata* | 4.00 | - | - | - | - |
| *Crithagra scotops* | 4.00 | 2.88 | 4.87 | 14.72 | -40.39 |

| | | | | | |
|---|---|---|---|---|---|
| *Crithagra sulphurata* | 4.00 | 3.39 | 6.13 | 10.13 | -20.58 |
| *Crithagra flaviventris* | 5.00 | 5.74 | 5.27 | 21.78 | -21.89 |
| *Crithagra flavigula* | 6.00 | 1.10 | 9.46 | 9.93 | - |
| *Crithagra dorsostriata* | 6.25 | 1.06 | 7.86 | 8.70 | -11.83 |
| *Crithagra mennelli* | 6.25 | 1.74 | 3.22 | 4.50 | -43.15 |
| *Crithagra mozambica* | 6.68 | 1.62 | 6.08 | 8.21 | -31.00 |
| *Crithagra whytii* | 6.68 | - | - | - | - |
| *Crithagra citrinipectus* | 7.50 | 3.36 | 7.67 | 14.89 | - |
| *Crithagra xantholaema* | 8.00 | 1.56 | 6.30 | 9.28 | -36.16 |

**Table S7.** Colour Elaboration Index and vocal trait values for each species included in the genus-level analysis of *Spinus*. CEI: Colour Elaboration Index; RI: Repertoire Index. Syllable rate and vocal deviation are expressed in Hz. Species with dashes in all their vocal traits could not be analysed due to the lack of high-quality recordings. Species with dashes only in the "Vocal deviation" column indicate songs that did not contain trills. Note that the column "Song length" is absent in this table because 10-second random fragments were used for the analysis.

| **Species** | **CEI** | **Syllable rate** | **RI** | **Vocal deviation** |
|---|---|---|---|---|
| *Spinus pinus* | 1.50 | 9.18 | 6.51 | -22.94 |
| *Spinus thibetanus* | 2.00 | 3.01 | 1.31 | -32.30 |
| *Spinus psaltria* | 3.00 | 5.65 | 3.58 | -28.91 |
| *Spinus xanthogastrus* | 3.99 | 3.41 | 5.66 | -23.28 |
| *Spinus atratus* | 4.00 | 4.72 | 6.79 | -27.85 |
| *Spinus uropygialis* | 4.50 | 4.27 | 0.94 | - |
| *Spinus spinus* | 5.00 | 3.52 | 8.64 | -25.82 |
| *Spinus tristis* | 5.00 | 3.13 | 2.57 | -23.60 |
| *Spinus atriceps* | 5.32 | 4.42 | 0.00 | -24.96 |
| *Spinus dominicensis* | 6.00 | - | - | - |
| *Spinus spinescens* | 6.68 | 4.84 | 0.00 | -24.31 |
| *Spinus crassirostris* | 6.68 | 4.99 | 1.31 | -29.21 |
| *Spinus magellanicus* | 6.68 | 4.62 | 5.13 | -24.37 |
| *Spinus barbatus* | 6.68 | 2.91 | 3.91 | -25.77 |
| *Spinus notatus* | 8.00 | 5.10 | 1.79 | -22.77 |
| *Spinus lawrencei* | 8.00 | 4.84 | 3.82 | -30.95 |
| *Spinus yarrellii* | 8.00 | - | - | - |
| *Spinus siemiradzkii* | 8.00 | - | - | - |
| *Spinus olivaceus* | 8.00 | 3.78 | 4.68 | -23.90 |
| *Spinus cucullatus* | 8.00 | 4.20 | 1.00 | -36.47 |
